# Microtubule inhibitors identified through non-biased screening enhance DNA transfection efficiency by delaying p62-dependent ubiquitin recruitment

**DOI:** 10.1101/2021.05.13.443985

**Authors:** Megumi Tsuchiya, Hidesato Ogawa, Kento Watanabe, Takako Koujin, Chie Mori, Kazuto Nunomura, Bangzhong Lin, Akiyoshi Tani, Yasushi Hiraoka, Tokuko Haraguchi

## Abstract

Ectopic gene expression is an indispensable tool in biology and medicine, but is often limited by the low efficiency of DNA transfection. We previously reported that depletion of the autophagy receptor p62/SQSTM1 enhances DNA transfection efficiency by preventing the degradation of transfected DNA. Therefore, p62 is a potential target for drugs to increase transfection efficiency. To identify such drugs, a non-biased high-throughput screening was applied to over 4,000 compounds from the Osaka University compound library, and their p62-dependency was evaluated. The top-scoring drugs were mostly microtubule inhibitors, such as colchicine and vinblastine, and all of them showed positive effects only in the presence of p62. To understand the p62-dependent mechanisms, the time required for p62-dependent ubiquitination, which is required for autophagosome formation, was examined using polystyrene beads that were introduced into cells as materials that mimicked transfected DNA. Microtubule inhibitors caused a delay in ubiquitination. Furthermore, the level of phosphorylated p62 at S405 was markedly decreased in the drug-treated cells. These results suggest that microtubule inhibitors inhibit p62-dependent autophagosome formation. Our findings demonstrate for the first time that microtubule inhibitors suppress p62 activation as a mechanism for increasing DNA transfection efficiency and provide solutions to increase efficiency.

## Introduction

Gene delivery is one of the most important steps in gene therapy and genetic modification in basic science. Gene therapy has great potential in clinical medicine, and this concept has become well established in therapeutic approaches. In basic science, DNA transfection is a powerful tool that enables the study of gene functions and their products in cells.

Endocytosis is a crucial pathway for the regulation of cellular uptake of plasmid DNA (Rejman *et al*. 2005; Khalil *et al*. 2006; Midoux *et al*. 2008). Importantly, endocytosis can promote the induction of selective autophagy, also called xenophagy (Chen *et al*. 2014). Generally, autophagy is a cytosolic bulk degradation pathway for the recycling of biomolecules through nonspecific degradation of proteins and organelles under nutrient starvation conditions. In contrast, selective autophagy plays an important defensive role against cellular infection by pathogens, as part of a starvation-independent autophagic defense system (Nakatogawa *et al*. 2009; Mizushima *et al*. 2011; Galluzzi *et al*. 2017). The conjugation of ubiquitin (Ub) to target pathogens is an important initial process in selective autophagy. Ubiquitination assists in the recruitment of autophagy receptor proteins, including p62/sequestosome-1 (p62/SQSTM1), against pathogens or transfected DNA (Dupont *et al*. 2010; Roberts *et al*. 2013; Chen *et al*. 2014; Alomairi *et al*. 2015; Tsuchiya *et al*. 2016). Therefore, suppression of the autophagy pathway can be a target for drugs to increase transfection efficiency.

We have previously reported that the depletion of the p62/SQSTM1 protein (hereafter designated p62) greatly increases the efficiency of DNA transfection in cultured cells (Tsuchiya *et al*. 2016). To monitor the behavior of the transfected DNA, we developed an experimental system using DNA-conjugated beads that mimic the transfected DNA. In this system, DNA-conjugated beads together with pHrodo, a fluorescent marker for endosome rupture, were incorporated into cells, and their intracellular dynamics were analyzed in a live cell (Tsuchiya *et al*. 2016). Using this system, we demonstrated that the transfected DNA was incorporated into cells through endocytosis, released from the endosomes into the cytosol, and entrapped via autophagy in a p62-dependent manner. Furthermore, we demonstrated that the recruitment of Ub around the transfected material was significantly delayed in p62 gene-knockout murine embryonic fibroblast (p62KO-MEF) cells compared with that in normal MEF cells (Tsuchiya *et al*. 2018). Additionally, p62 phosphorylation at S405 (equivalent to human S403) is a crucial step for the recruitment of Ub to the target site of transfected materials that mimic ectopic DNA (Tsuchiya *et al*. 2016). Hence, p62 phosphorylation at S405 is an essential step for transfection-induced selective autophagy (Tsuchiya *et al*. 2018). As p62 plays an important role in gene delivery through initial ubiquitination of the transfected DNA, inhibition of p62 may increase transfection efficiency. The aim of this study was to identify a small chemical compound that blocks the initial step of p62-dependent selective autophagy to enhance DNA transfection efficiency in mammalian cells.

## Results

### High-throughput screening for drugs enhancing transfection efficiency

To identify potential compounds that can enhance transfection efficiency, high-throughput screening based on a luciferase assay was performed on MEF cells using an automated workstation (Fig. 1A). The MEF cells were seeded in 384-well plates and incubated for 18 h with each compound from the Osaka University compound library (4,400 compounds; Tachibana *et al*. 2018) at a final concentration of 10 µM. The negative control was dimethyl sulfoxide (DMSO) at a final concentration of 1%. The cells were transfected with the pCMV-Luc plasmid (encoding luciferase driven by the cytomegalovirus immediate early (CMV-IE) promoter) for 28 h using Lipofectamine 2000. The cells were subjected to a cell viability assay, followed by a luciferase assay (Fig. 1A). Luciferase activity, as expressed from the transfected plasmid, was measured, and transfection efficiency was evaluated based on luciferase activity normalized to cell viability. Among the 4,400 compounds tested, 160 were removed from the list for selection as valid compounds because of severe defects in cell morphology. The transfection efficiency of the cells treated with each of the remaining 4,240 compounds is plotted in Fig. 1B. In this first screening, out of the 4,240 tested compounds, we identified 87 compounds that increased luciferase activity compared with that in the negative control (approximately 2.1% positive hit rate). The cutoff value used for selection was the mean value of the DMSO control + 4 × standard deviation (SD) (mean 0.347, SD = 0.242).

**Fig. 1.**
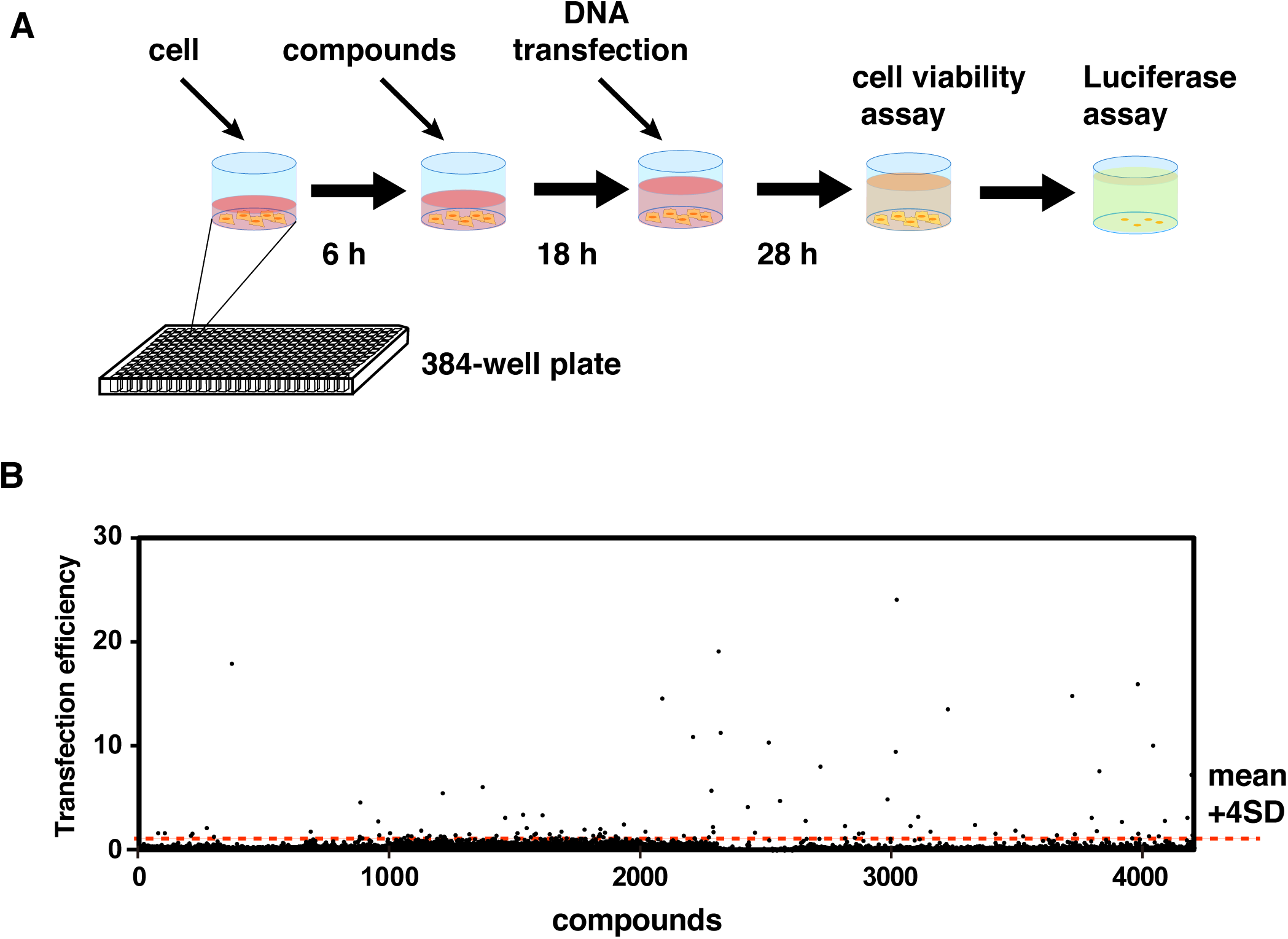
High-throughput screening for drugs enhancing transfection efficiency. (A) Schematic diagram of the process of high-throughput screening using automated workstations. Murine embryonic fibroblast (MEF) cells were treated with each compound and then transfected with the pCMV-Luc plasmid in the presence of each compound. Cell viability and the luciferase activity from the transfected plasmid were measured. Transfection efficiency was evaluated based on luciferase activity normalized to cell viability. (B) High-throughput screening was performed for 4,400 compounds from the Osaka University compound library as described. Values of transfection efficiency were plotted for the selected 4,240 compounds. Dimethyl sulfoxide (DMSO) was used as the negative control. The red broken line represents the mean + 4 × SD of all negative control assay points.

The activities of these 87 compounds were further analyzed via a second screening. To achieve this, MEF cells were cultured in 96-well plates and transfected with the pCMV-Luc plasmid for 28 h using Lipofectamine 2000 in the presence of each of these compounds at a concentration of 1 μM (Fig. 2A). Transfection efficiency was determined by a luciferase assay, in which luciferase activity expressed from the transfected plasmid was measured and normalized to cell viability, as described in Experimental procedures. The transfection efficiency values for the 87 compounds are listed in Table S1. Notably, the top 9 compounds with high transfection efficiencies (> 20-fold that of the negative control) were all microtubule inhibitors (Fig. 2B; Table S1).

**Fig. 2.**
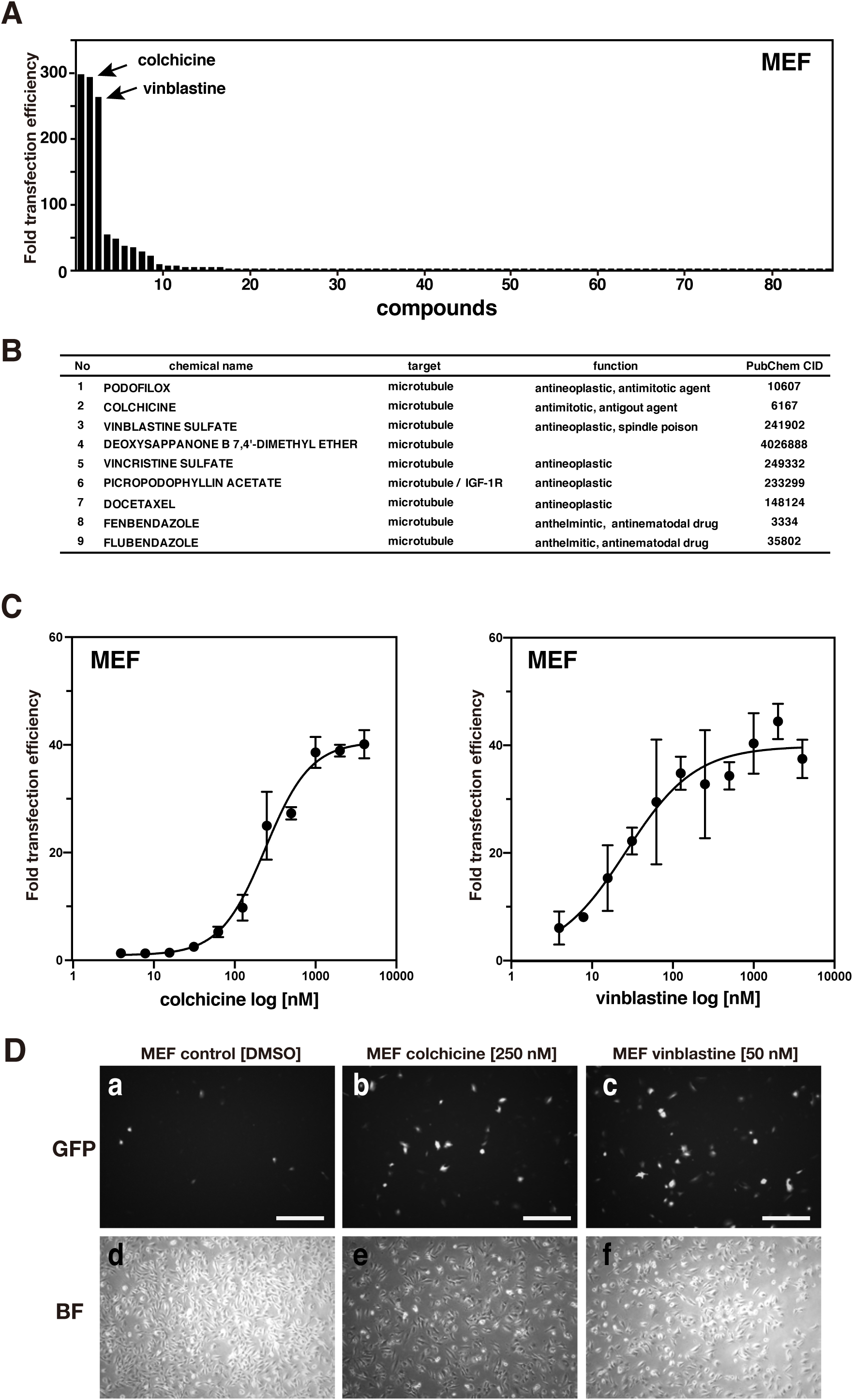
Second screening of compounds selected through high-throughput screening. (A) The top 87 compounds identified in the primary screen were further examined. MEF cells were transfected with the pCMV-Luc plasmid for 28 h in the presence of 1 μM of each compound. DMSO was used as the negative control. Transfection efficiency was determined based on luciferase activity normalized to the cell viability. Bar graphs show fold changes in transfection efficiency relative to the value from the control DMSO-treated cells in descending order. The second and third columns represent data for colchicine and vinblastine as indicated. (B) The top 9 potential compounds for enhancement of transfection efficiency. (C) MEF cells were transfected with the pCMV-Luc plasmid for 28 h in the presence of colchicine (left graph) or vinblastine (right graph) at the indicated concentrations. Transfection efficiency was determined as described above. Filled circles and vertical lines indicate the mean and SD, respectively, of at least three independent experiments. (D) Control DMSO-treated MEF cells (a, d), 250 nM colchicine-treated MEF cells (b, e), and 50 nM vinblastine-treated MEF cells (c, f) were examined using fluorescence microscopy 24 h after transfection with a GFP-expressing plasmid pCMX-AFAP (upper panels; GFP); the lower panels represent the corresponding bright-field (BF) images of the upper panels. Scale bar = 500 μm.

As microtubule inhibitors were found to increase transfection efficiency, we also tested nocodazole, an extensively used microtubule inhibitor, which was not included in the Osaka University compound library. Transfection efficiency was determined in the presence or absence of 1 μM nocodazole in 96-well plates (Fig. S1). The results show that nocodazole has a transfection-enhancing activity at 1 μM, similar to colchicine (Fig. S1).

We further examined whether microtubule inhibitors are effective against various transfection reagents with different mechanisms. Effectene (a non-liposomal lipid reagent) and Viofectin (a non-lipid high-charge cationic polymer reagent) were examined in addition to the previously used Lipofectamine 2000 (a cationic lipid reagent). In the presence of 300 nM colchicine, the transfection efficiencies in all these transfection reagents increased, although the degree of effectiveness varied (Fig. S2). The increase in transfection efficiency in the presence of 300 nM colchicine was approximately 2-, 6-, and 29-fold in Effectene, Viofectin, and Lipofectamine 2000, respectively (Fig. S2), suggesting that colchicine is effective with various non-viral transfection reagents.

### Microtubule inhibitors enhance gene transfection efficiency

To further evaluate the effects of microtubule inhibitors on DNA transfection efficiency, we selected two commonly used microtubule inhibitors, colchicine and vinblastine. These were ranked second and third, respectively, in the 2^nd^ screening (Fig. 2A). First, we examined the dose-dependency of these inhibitors in MEF cells (Fig. 2C). Cells were treated with colchicine or vinblastine at various concentrations and transfected with the pCMV-Luc DNA plasmid. Then, their luciferase activity was measured to determine transfection efficiency. Transfection efficiency increased in a dose-dependent manner in the MEF cells (Fig. 2C). Statistical analysis showed that the EC50 (half-maximal effective concentration) of colchicine and vinblastine was 239.1 and 26.29 nM, respectively, in the MEF cells (Fig. 2C). Furthermore, we evaluated transfection efficiency by observing GFP expression in MEF cells. The number of GFP-positive cells was greater in colchicine- and vinblastine-treated cells than in DMSO-treated control cells, and cell viability was approximately 70% in both drug-treated cells compared with that in the control under these conditions (Fig. 2D). These results suggest that treatment with colchicine and vinblastine can enhance transfection efficiency.

As microtubule depolymerization activates the transcription factor NF-κB to induce NF-κB-dependent gene expression (Rosette & Karin, 1995) and the CMV promoter in the pCMV-Luc plasmid possesses an NF-κB binding site (Rosette & Karin, 1995; Wang & MacDonald, 2004), treatment with microtubule inhibitors may induce luciferase gene expression through NF-κB-mediated CMV promoter activation. To eliminate this possibility, we tested promoter activity in a cell line (MEF-LUC cells) with a CMV-driven luciferase plasmid integrated into the genome. In these cells, we measured the levels of luciferase activity in the presence or absence of colchicine or vinblastine. Treatment with colchicine for 27 h increased luciferase gene expression only marginally (approximately 1.2-fold), which did not explain the considerably greater increase (approximately 40-fold) in the transfected DNA plasmids (Fig. S3A; compare with Fig. 2C). Treatment with vinblastine also yielded a similar result, with almost no increase in luciferase expression (Fig. S3B; compare with Fig. 2C). These results indicate that the increase in transfection efficiency in the presence of microtubule inhibitors does not reflect NF-κB-dependent promoter activation.

### Transfection enhancement occurs in the presence of autophagy receptor p62

As p62 acts as an inhibitory factor for DNA transfection by recruiting LC3 (microtubule-associated protein 1A/1B-light chain 3; also called ATG8), a marker protein for autophagosomes, to transfected DNA (Tsuchiya *et al*. 2016), the transfection-enhancing activity of the tested compounds may be deduced by the invalidation of the p62-dependent autophagic pathway. Based on this hypothesis, we examined the activity of the 87 compounds at a concentration of 1 μM in p62KO-MEF cells under the same conditions as those in Fig. 2A (Fig. 3A). All compounds exhibited little or no transfection-enhancing activity in p62KO-MEF cells (Fig. 3A). As the microtubule inhibitors (compounds 1–9 in Fig. 2B) seemed to show a slight increase in transfection enhancing activity (Fig. 3A), we further tested colchicine and vinblastine for transfection-enhancing activity at various concentrations from 15.6 to 2000 nM (Fig. 3B, C) and found that these two microtubule inhibitors did not show transfection-enhancing activity at any of the tested concentrations (Fig. 3B, C). This result suggests that microtubule inhibitors function in a p62-dependent autophagic pathway.

**Fig. 3.**
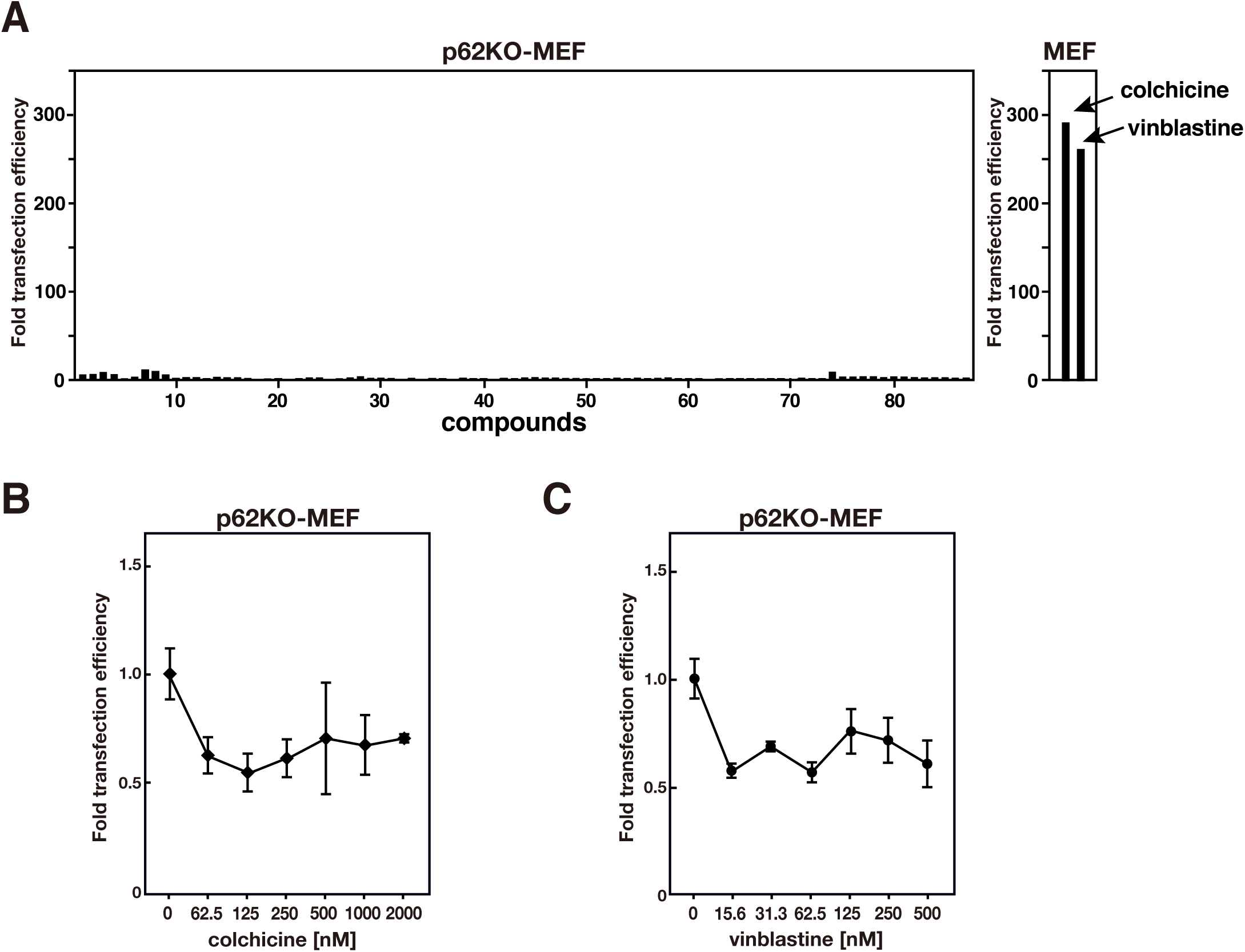
Colchicine- and vinblastine-treated MEF cells do not enhance transfection efficiency in the absence of p62. (A) The top 87 compounds were examined in p62KO-MEF cells. The p62KO-MEF cells were transfected with the pCMV-Luc plasmid for 28 h in the presence of 1 μM of each compound, and transfection efficiency was determined as described in the legend to Fig. 2A. The fold changes in transfection efficiency relative to the value from the control DMSO-treated cells are shown in the same order as in Fig. 2A. The two columns on the right are reproduced from Fig. 2A for comparison. (B, C) The p62KO-MEF cells were transfected for 28 h with the pCMV-Luc plasmid in the presence of the indicated concentrations of colchicine (B) or vinblastine (C). Each value of the transfection efficiency is indicated as the mean ± SD of at least three independent experimental results.

### p62-dependent ubiquitination is delayed by microtubule inhibitors

To understand the molecular mechanisms of the p62-dependent enhancement of transfection efficiency by colchicine and vinblastine, we employed an experimental method using polystyrene beads that had been developed to monitor the behavior of the transfected DNA (Kobayashi *et al*. 2010) (Fig. 4A). In this method, the beads are incorporated into cells with transfection reagents via endocytosis and enter the cytosol after rupture of the endosomal membrane. The beads that appeared in the cytosol were targeted for autophagy, similar to the transfected DNA (Kobayashi *et al*. 2010; Kobayashi *et al*. 2015) (Fig. 4A). For monitoring in living cells, the beads were pre-conjugated with pHrodo dye, which emits fluorescence under acidic pH conditions, such as those in the acidic endosome but not in the cytosol. Therefore, this dye serves as a marker of endosome membrane rupture, as previously described (Kobayashi *et al*. 2010). The pHrodo-conjugated beads were incorporated into MEF cells expressing GFP-fused Ub protein (GFP-Ub MEF cells) (Tsuchiya *et al*. 2018), and the assembly of GFP-Ub around the beads was observed in a living cell using time-lapse fluorescence microscopy (Fig. 4B). In the control DMSO-treated cells, the time for Ub recruitment to the beads was approximately 3–4 min (Fig. 4B, middle panel) after the disappearance of pHrodo fluorescence (Fig. 4B, upper panel). In contrast, the time for GFP-signal accumulation was 9–10 min in 500 nM colchicine-treated MEF cells (Fig. 4C, middle panel), which was longer than that in the control cells (Fig. 4B). Statistical analysis was performed to determine the timing of GFP-signal accumulation around the beads after the loss of pHrodo signals in cells expressing GFP-Ub with or without the inhibitor (Fig. 4D). In the DMSO-treated cells, the time for Ub recruitment to the beads was ∼4 min (median) after the disappearance of pHrodo fluorescence (mean and SD, 4.167 ± 1.88 min, *n* = 24 beads; lane 1 in Fig. 4D). Moreover, the timing of GFP-signal accumulation was ∼6 min (median) in the 100 nM colchicine-treated MEF cells (mean and SD: 8.286 ± 8.36 min, *n* = 28 beads: lane 2 in Fig. 4D), ∼11 min (median) in the 500 nM colchicine-treated MEF cells (mean and SD: 11.32 ± 11.30 min, *n* = 22 beads: lane 3 in Fig. 4D), and ∼6 min (median) in the vinblastine-treated MEF cells (mean and SD: 6.48 ± 3.81 min, *n* = 31 beads; lane 4 in Fig. 4D). Additionally, the GFP-Ub signals did not accumulate within 60 min in colchicine- or vinblastine-treated GFP-Ub MEF cells (these beads were not counted and were not included in the *n*; upper column in Fig. 4D). These results show that GFP-Ub accumulation around the beads was significantly delayed in colchicine- or vinblastine-treated MEF cells. Colchicine and vinblastine have different effects on microtubule depolymerization; colchicine forms diffusely distributed tubulins/microtubules, whereas vinblastine forms the paracrystal structure of the microtubules (Haraguchi *et al*. 1997). Nonetheless, both have the same function of increasing DNA transfection efficiency, suggesting that normal microtubule function or structure is important for Ub recruitment to the target site.

**Fig 4.**
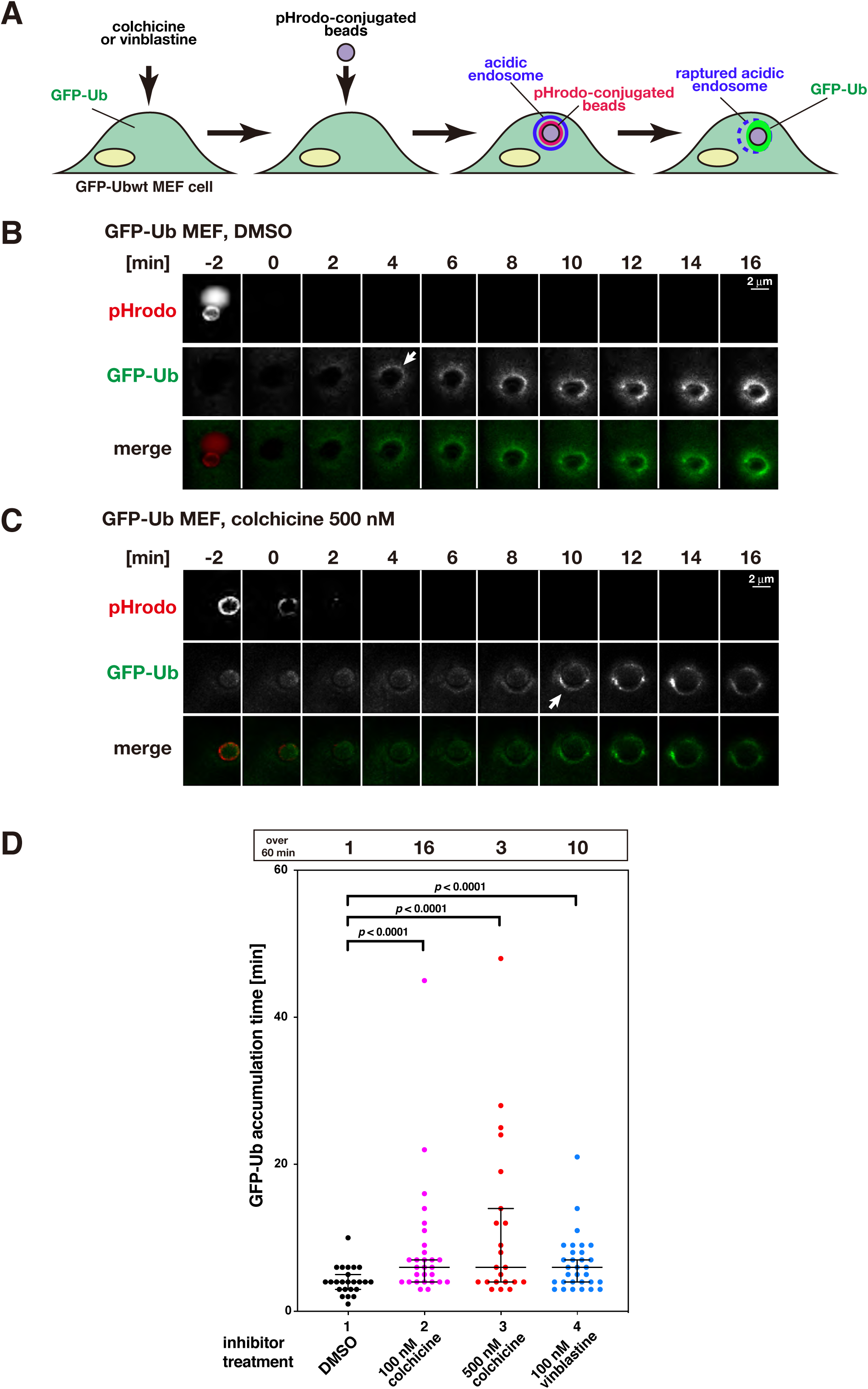
Colchicine- and vinblastine-treated MEF cells affect the timing of ubiquitination. (A) Schematic diagram of the experimental system using beads incorporated into cells to monitor the behavior of transfected materials. (B, C) Time-lapse images of pHrodo and GFP-Ub fluorescence around a single pHrodo-conjugated bead in MEF cells. Images were obtained every minute for approximately 60 min. The panels show representative images of pHrodo and GFP-Ub fluorescence. (B) GFP-Ub MEF cells treated with DMSO as a control. Arrows indicate the timing of appearance of GFP-Ub signals around the beads. (C) GFP-Ub MEF cells treated with 500 nM colchicine. Arrows indicate the timing of appearance of GFP-Ub signals around the beads. Scale bar = 2 μm. (D) Statistical analysis was performed for the time taken for GFP-signal accumulation around the beads after the loss of pHrodo signals in the GFP-Ub MEF cells. Lane 1: control treated with DMSO. Lane 2: 100 nM colchicine-treated MEF cells. Lane 3: 500 nM colchicine-treated MEF cells. Lane 4: 100 nM vinblastine-treated MEF cells. The median values were 4 min for GFP-Ub (*n* = 24 beads), 6 min for 100 nM colchicine (*n* = 28 beads), 6 min for 500 nM colchicine (*n* = 22 beads), and 6 min for 100 nM vinblastine (*n* = 31 beads). Three independent experiments were performed for each lane, and the total bead number is indicated as *n*. Statistical differences (*p* < 0.0001) were determined using the Kruskal–Wallis test. Error bars indicate 95% confidence intervals.

### Active form of p62 is suppressed by microtubule inhibitors

It has been reported that the phosphorylated form of p62 at amino acid residue S405 (p62 S405) is required for Ub recruitment during selective autophagy (Tsuchiya *et al*. 2018). Therefore, Ub recruitment can be delayed by a decrease in the level of phosphorylated p62 at S405. To test this hypothesis, we performed Western blot analysis to evaluate the total p62 protein levels and p62 S405 phosphorylation levels (Fig. 5). The p62 S405 phosphorylation levels were very low before DNA transfection (“no transfection” in Fig. 5). However, after transfection (“DNA transfection 24 h”), these levels greatly increased in DMSO-treated MEF cells, although the p62 levels remained unchanged. This suggests that DNA transfection induced an increase in p62 S405 phosphorylation. However, in MEF cells treated with colchicine or vinblastine, the level of p62 S405 phosphorylation decreased. This suggests that microtubule inhibitors inhibit Ub recruitment by decreasing the level of phosphorylated p62 at S405.

**Fig. 5.**
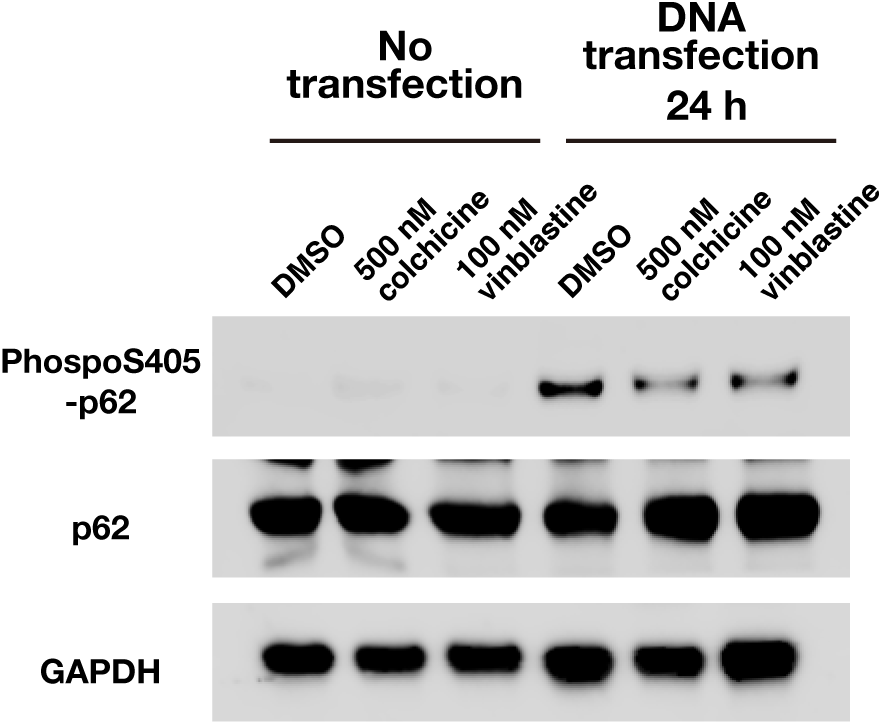
Colchicine- and vinblastine-treated MEF cells decrease the level of phosphorylated p62 at S405. Western blot analysis was performed for total p62 and S405-phosphorylated p62 in MEF cells. Cells were pretreated with 500 nM colchicine or 100 nM vinblastine for 30 min, and then transfected with the pCMX-AFAP plasmid in the presence of 500 nM colchicine or 100 nM vinblastine. DMSO was used as the control. Samples were subjected to Western blot analysis with no transfection or with transfection for 24 h. GAPDH was used as the loading control.

## Discussion

### Microtubule structure/function and transfection efficiency

In this study, we used a non-biased, high-throughput screening approach to identify small chemical compounds that increase DNA transfection efficiency and identified microtubule inhibitors as the top 9 compounds. Therefore, inhibition of the microtubule structure or function seems to trigger an increase in transfection efficiency. This finding is consistent with previous reports that microtubule depolymerizing reagents, nocodazole, increased transfection efficiency in CV-1 cells (Lindberg *et al*. 2001) and colchicine and vinblastine increased transfection efficiency in cultured vascular smooth muscle cells (Wang & MacDonald 2004). Additionally, microtubule-polymerizing reagents, such as paclitaxel, also increase the transfection efficiency in COS-7 and A549 cells (Hasegawa *et al*. 2001; Nair *et al*. 2002). The polymerizing agent docetaxel, a chemical compound closely related to paclitaxel, was included in our screening results for the top 9 compounds (Fig. 2B, Rank 7). Moreover, tubulin deacetylation inhibitors, such as HDAC6 inhibitors, also increase transfection efficiency in A549 cells, PC3 cells, and mesenchymal stem cells (Vaughan *et al*. 2008; Barua & Rege 2010; Ho *et al*. 2017). This is likely because acetylated tubulin can stabilize microtubule structures, which in turn implies that microtubule structure stabilization can also increase gene transfection efficiency. Taken together, these findings strongly suggest that inhibition of the intact structure or dynamic nature of microtubules is important for increasing the efficiency of gene transfection.

### Microtubule inhibitors affect selective autophagy pathways

Our results showed that treatment of cells with colchicine or vinblastine increased transfection efficiency in a p62-dependent manner. However, it is difficult to attribute these observations to a particular cause. Microtubules are major contributors to the trafficking of several components in the endosomal/lysosomal pathway; hence, it is logical to propose that these inhibitors may block the autophagy pathway. The role of microtubules in autophagosome formation appears to be different based on the culture medium conditions, e.g., vegetatively growing medium (basal) or starvation medium (inducible) conditions. Several studies have used microtubule inhibitors under basal conditions to show that microtubules do not participate in autophagosome formation (Reunanen *et al*. 1988; Aplin *et al*. 1992; Kochl *et al*. 2006). However, under inducible conditions, disassembling the microtubules with these inhibitors prevented autophagosome formation, suggesting that microtubules play a crucial role in this process (Kochl *et al*. 2006; Geeraert *et al*. 2010). Previous studies have shown that transfected DNA also induces the selective autophagy pathway (Roberts *et al*. 2013; Chen *et al*. 2014; Tsuchiya *et al*. 2016); hence, microtubule inhibitors may affect transfection-induced autophagosome formation. The detailed mechanisms of the inhibition of autophagosome formation following microtubule inhibitor treatment are not clear; however, several autophagy factors are associated with microtubules, including ATG8 (LC3), ATG1 (ULK1), ATG6 (Beclin1), ATG18 (WIPI1), and p62 (Mackeh *et al*. 2013). LC3 has long been thought to be involved in the regulation of microtubule assembly and disassembly (Mann & Hammarback 1994; Deretic 2008). Furthermore, ATG18-positive pre-autophagosomal structures can move along microtubules, and this movement is extremely sensitive to microtubule inhibitor treatment (Polson *et al*. 2010). These data suggest that microtubules contribute to the sequestration, recruitment, and movement of autophagy factors for the formation of inducible autophagosomes. Therefore, microtubule inhibitors may block the inducible autophagy pathway, resulting in a decrease in the likelihood of DNA degradation and increase in transfection efficiency.

### Microtubule inhibitors decrease the level of phosphorylated p62

The ubiquitination of endosome membrane proteins surrounding exogenous material, such as transfected DNA, is the initial step in inducible selective autophagy (Randow 2011; Wileman 2013; Tsuchiya *et al*. 2016). In our study, treatment with colchicine or vinblastine delayed the recruitment of Ub proteins. This delay in recruitment is also caused by the depletion of p62 or mutation of the p62 phosphorylation site at S405 (human S403) (Tsuchiya *et al*. 2018). As p62 has been reported as a microtubule-associated factor (Geeraert *et al*. 2010), treatment with a microtubule inhibitor may affect p62-mediated Ub recruitment. Specifically, our results showed that p62 S405 phosphorylation was significantly impaired by treatment with colchicine or vinblastine after transfection. p62 S405 is phosphorylated by kinases such as CK2 and TBK1 (Matsumoto *et al*. 2011; Matsumoto *et al*. 2015). These kinases are also associated with tubulin (Faust *et al*. 1999; Lim *et al*. 2004; Pillai *et al*. 2015; Yang *et al*. 2018). After transfection, the recruitment of these kinases may be affected by treatment with microtubule inhibitors. Further studies are necessary to elucidate the detailed mechanisms by which these microtubule inhibitors affect the phosphorylation levels of p62.

In this study, a non-biased high-throughput screening method revealed that microtubule inhibitors are the most effective reagents for increasing transfection efficiency among the drugs present in the Osaka University compound library, and revealed for the first time, the mechanism by which microtubule inhibitors increase transfection efficiency; that is, microtubule inhibitors suppress p62 phosphorylation, which is required for autophagosome formation to degrade transfected DNA. Our findings provide solutions to increase transfection efficiency.

## Experimental procedures

### Plasmids

pCMV-Luc plasmid encoding the luciferase gene *luc2*, used as a luciferase expression vector in this study, was purchased from Promega (E1310; pGL4.50 [*luc2*/CMV/Hygro] vector; Promega, Madison, WI, USA). The GFP expression plasmid pCMX-AFAP was prepared as previously described (Ogawa *et al*. 2004).

To create cells stably expressing luciferase (see the next section), we constructed the PB-CMV-LUC-Zeo vector. We first constructed the PB-EF1-MCS-IRES-Zeo vector. To insert the zeocin resistance gene DNA sequence, two DNA fragments (#1 and #2) were amplified as follows: the cloning site with Kozak sequence fragment #1 was amplified from the PB-EF1-MCS-IRES-Neo vector (PB533A-2; System Biosciences, Palo Alto, CA, USA) using PCR and the following primers: 5′-CTGAAGGATGCCCAGAAGGTACCCCATTGT-3′ and 5′-TCCGGACGCCATGGTTGTGG-3′. The zeocin coding fragment #2 was amplified from the pcDNA3.1/Zeo (+) vector (V86020; Thermo Fisher Scientific, Yokohama, Japan) using PCR and the following primers: 5′-ACAACCATGGCGTCCGGAATGGCCAAGTTGACCAGTGCCGTTCC-3′ and 5′-TCCAGAGGTTGATTGTCGACTCAGTCCTGCTCCTCGGCCACGAA-3′. Following digestion with KpnI and SalI, fragments #1 and #2 were inserted into the PB-EF1-MCS-IRES-Neo vector using the In-Fusion HD Cloning Kit (639648; Takara Bio Inc. Kusatsu, Japan). This resulted in the PB-EF1-MCS-IRES-Zeo vector. Next, the human cytomegalovirus immediate early enhancer and promoter (CMV-IE) DNA sequence (#3) was amplified from the pGL4.50 [*luc2*/CMV/Hygro] vector using PCR and the following primers: 5′-GGGGATACGGGGAAAAGGCCTCGTTACATAACTTACGGTAAATG-3′ and 5′-GAATTCGCTAGCTCTAGAAGCTCTGCTTATATAGACCTCCCACC-3′. The luciferase DNA sequence (#4) was amplified from the pGL4.50 [*luc2*/CMV/Hygro] vector using PCR and the following primers: 5′-TCTAGAGCTAGCGAATTCATGGAAGATGCCAAAAACATTAAGAA-3′ and 5′-CGATTTAAATTCGAATTCTTACACGGCGATCTTGCCGCCCTTCT-3′. Following digestion with EcoRI and StuI, fragments #3 and #4 were inserted into the PB-EF1-MCS-IRES-Zeo vector using the In-Fusion HD Cloning Kit. This resulted in the PB-CMV-LUC-Zeo vector.

### Cell strains

p62KO-MEF cells (p62^-/-^ cells) and their parental MEF cells were kindly provided by Dr. Tetsuro Ishii (University of Tsukuba) (Komatsu *et al*. 2007). MEF cells stably expressing GFP-Ub were generated as previously described (Tsuchiya *et al*. 2018). To obtain MEF cells or p62KO-MEF cells stably expressing luciferase, MEF cells or p62KO-MEF cells were transfected with the PB-CMV-LUC-Zeo plasmid (generated as described in the previous section) and cultured in the presence of 100 μg/mL Zeocin (R25501, Thermo Fisher Scientific), and then, single clones (MEF-LUC cells) were selected. Each stable clone was examined for luciferase protein expression using a luciferase assay.

### Cell culture

All cell lines were maintained in culture medium (Dulbecco’s modified Eagle medium (DMEM) (D6429; Sigma-Aldrich, St. Louis, MO, USA) supplemented with 10% fetal bovine serum) in the presence of 5% CO_2_ at 37°C.

### High-throughput screening for enhancer compounds

The MEF cells were seeded at a density of 0.8 × 10^3^ cells per well (384-well microplate, 781091; Greiner Bio-One, Tokyo, Japan) using a Multidrop COMBI (Thermo Fisher Scientific) and cultured for 6 h in culture medium. The cells were pretreated for 18 h with 1% DMSO (negative control) or the screening compounds (10 μM each) using a Fluent780® Automation Workstation (Tecan Japan, Kawasaki, Japan) with a 96-channel head adapter and Tecan sterile tips (30048824; Tecan Japan). The pretreated cells in each well were transfected with 25 ng of pCMV-Luc plasmid using Lipofectamine 2000 (Thermo Fisher Scientific) for 28 h in the presence of each compound. Cell viability was measured with the RealTime-Glo™ MT Cell Viability Assay (E9713; Promega) using the GloMax® Discover Microplate Reader (Promega). Luciferase activity was measured with the ONE-Glo™ Luciferase Assay System (E6120; Promega) using the GloMax® Discover Microplate Reader according to the manufacturer’s protocol. The values of luciferase activity were normalized to the cell number obtained from cell viability measurements.

### Transfection efficiency measurements

Transfection efficiency was determined based on the luciferase activity expressed from the transfected plasmid. Cells were seeded at a density of 0.45 × 10^4^ cells per well (96-well assay plate; 3603 Corning, Corning, NY, USA) and cultured overnight in culture medium. The cells were pretreated with 1% DMSO (negative control) or the screening compounds (0.1–10 μM each) for 30 min. The cells were then transfected with 100 ng of pCMV-Luc plasmid using Lipofectamine 2000 for 28 h in the presence of each compound. Cell viability was measured with the RealTime-Glo™ MT Cell Viability Assay using the GloMax® 96 Microplate Luminometer (Promega). Luciferase activity was measured with the ONE-Glo™ Luciferase Assay System using the GloMax® 96 Microplate Luminometer according to the manufacturer’s protocol. The values of luciferase activity were normalized to the cell number obtained from cell viability measurements. Mean EC50 values and SDs were determined from three independent experiments. Transfection efficiencies were determined by luciferase activity normalized to cell viability.

Viofectin (VFT1001; Viogene, Nacalai tesque, Kyoto, Japan) and Effectene (301425; Qiagen, Tokyo, Japan) were also used as transfection reagents. For Viofectin, the same transfection procedure as that described above for Lipofectamine 2000 was used, except that the reagents were different. For Effectene, some procedures were modified as follows: initial cell density was 1 × 10^4^ cells per well and the cells were transfected for 4 h and further incubated for another 20 h in culture medium without transfection reagents.

### GFP expression

MEF cells were seeded at a density of 1.5 × 10^5^ cells per well (6-well assay plate; 3335 Corning, Corning, NY, USA) and cultured overnight in culture medium. The cells were pretreated with 1% DMSO (negative control), 250 nM cholchicine, or 50 nM vinblastine for 30 min. The cells were then transfected with 2 μg of pCMV-Luc plasmid using Lipofectamine 2000 for 24 h in the presence of each drug. The cells were observed using an objective lens (N Plan 5x/0.12 PH 0) on a Leica DM IL LED fluorescence microscope (Leica Microsystems, Wetzlar, Germany).

### Preparation of pHrodo-conjugated beads

pHrodo-conjugated beads were prepared as previously described (Kobayashi *et al*. 2010). Briefly, Dynabeads M-270 Streptavidin (DB65306; Thermo Fisher Scientific) were washed three times with phosphate buffered saline (PBS) and resuspended in 100 mM sodium bicarbonate buffer (pH 8.5) to an appropriate concentration (typically a 1:10 or 1:20 dilution). pHrodo-succinimidyl ester (P36600; Thermo Fisher Scientific) was then added to the bead suspension and incubated in sodium bicarbonate buffer for 1 h at room temperature (about 26°C). After the conjugation reaction, the beads were washed with sodium bicarbonate buffer and suspended in PBS.

### Incorporation of beads into living cells

One day before incorporating the beads, GFP-Ub MEF cells were seeded onto 35 mm glass-bottom culture dishes (P35G-1.5-10-C; MatTek, Ashland, MA, USA) at a density of 1.5 × 10^5^ cells/dish in culture medium. The cells were pretreated with each microtubule inhibitor (100 nM colchicine, 500 nM colchicine, or 100 nM vinblastine) or DMSO as a control for 30 min at 37°C in a CO_2_ incubator before bead incorporation.

The transfection reagent Effectene (301425; Qiagen, Tokyo, Japan) was used to incorporate the beads. Transfection-reagent-coated beads were prepared by mixing pHrodo-conjugated beads with the Effectene transfection reagent according to the manufacturer’s protocol, except that a bead suspension was used instead of the DNA solution. The resulting bead mixture (∼10 μL) was mixed with 90 μL of the culture medium containing each microtubule inhibitor and added to the cells by replacing the medium. After incubation for 1 h at 37°C in a CO_2_ incubator, the cells were washed twice with fresh growth medium to remove unattached beads and then further incubated with the medium containing each microtubule inhibitor or control DMSO for the time indicated in each experiment.

### Time-lapse imaging

Cells were treated with 100 ng/mL Hoechst 33342 (B2261; Sigma-Aldrich) for 15 min to stain chromosomes, as previously described (Haraguchi *et al*. 1997), except that the medium containing each microtubule inhibitor or control DMSO was used. After washing with the culture medium, the cells were cultured in fresh culture medium (phenol red-free DMEM containing 10% fetal bovine serum) containing each microtubule inhibitor or control DMSO. Time-lapse observations were performed using an oil-immersion objective lens (UApo40/NA1.35; Olympus, Tokyo, Japan) on a DeltaVision microscope system (GE Healthcare Life Sciences Japan, Tokyo, Japan) placed in a temperature-controlled room (37°C), as previously described (Haraguchi *et al*. 1997). Images were obtained every minute for approximately 60 min.

The timing of GFP-Ub fluorescence signal accumulation around the beads was determined as previously described (Tsuchiya *et al*. 2018). Briefly, the fluorescence intensity of the region (18 pixels square) surrounding the beads was measured using the FUJI software suite (IMAGE J; National Institutes of Health, Bethesda, MD, USA), and the background fluorescence intensity of the region with no beads in the same cell was subtracted.

### Western blot analysis

MEF cells were seeded at a density of 1.5 × 10^5^ cells per well (6-well assay plate; 3335 Corning, Corning, NY, USA) and cultured overnight in culture medium. The cells were pretreated with 1% DMSO (negative control), 500 nM colchicine, or 100 nM vinblastine for 30 min. The cells were then transfected with 2 μg of pCMX-AFAP plasmid using Lipofectamine 2000 for 24 h in the presence of each drug. Western blot analysis was performed as previously described (Tsuchiya *et al*. 2018). Briefly, cell lysates were prepared in a lysis buffer [50 mM Tris-HCl, pH 7.5, 150 mM NaCl, 1 mM EDTA, 1% Triton X-100, 1 x Phosphatase Inhibitor Cocktail Solution II (160-24371; FUJIFILM Wako Pure Chemical Corporation, Osaka, Japan) and 1 x protease inhibitor cocktail (Nacalai tesque Inc., Kyoto, Japan)]. The lysates were subjected to electrophoresis on NuPAGE 4% to 12% Bis-Tris gels (NP0321; Thermo Fisher Scientific). Proteins were transferred to polyvinylidene fluoride membranes and probed using anti-p62 (SQSTM1) (PM045; MBL, Nagoya, Japan), anti-Phospho-SQSTM1/p62(Ser403) (D8D6T; Cell Signaling Technology, Danvers, MA, USA), and anti-glyceraldehyde 3-phosphate dehydrogenase (GAPDH) (14C10; Cell Signaling Technology) antibodies, and secondary antibody conjugated to horseradish peroxidase (NA9340V; GE Healthcare Life Sciences). Protein bands were stained with ImmunoStar Zeta (295-72404; FUJIFILM Wako Pure Chemical Corporation) and detected by chemiluminescence using a ChemiDoc MP imaging system (Bio-Rad, Tokyo, Japan).

### Statistical analysis

The *p*-values were obtained by performing Kruskal-Wallis tests using GraphPad Prism 8 software (GraphPad Software, Inc., La Jolla, CA, USA).

## Acknowledgments

This work is dedicated to the late Professor Tomoko Ogawa. We are grateful to Dr. Eiji Warabi and Dr. Tetsuro Ishii (Tsukuba University) for providing p62KO-MEF cells and the corresponding wild-type MEF cells. We thank J. Iacona, Ph.D., from Edanz Group (https://en-author-services.edanzgroup.com/ac) for English-editing a draft of this manuscript. This study was supported by The JSPS Kakenhi Grant Numbers JP19K06488 to MT, JP19KK0218, JP20H05322 and JP20K07029 to HO, JP18H05533 and JP20H00454 to YH, and JP17K19505 and JP18H05528 to TH. This research was supported by Platform Project for Supporting Drug Discovery and Life Science Research (Basis for Supporting Innovative Drug Discovery and Life Science Research (BINDS)) from AMED under Grant Number JP20am0101084.

## Author contributions

MT, HO, KW, TK, CM, KN, BL and AT performed the experiments. MT, HO, AT, YH, and TH designed the experiments. All authors analyzed and discussed the data, and MT, HO, YH, and TH wrote the manuscript.

## Supporting Information

**Table S1.**
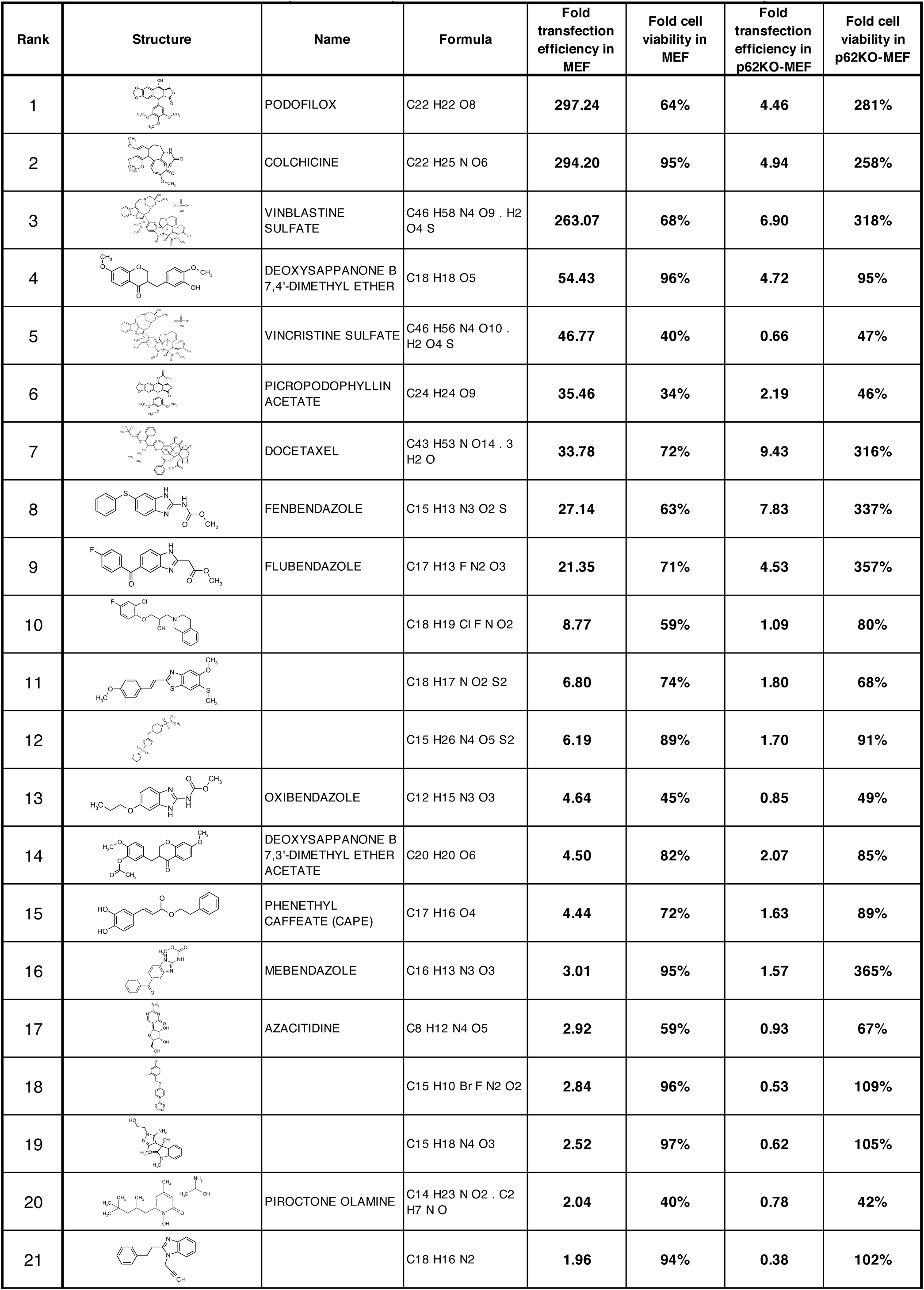
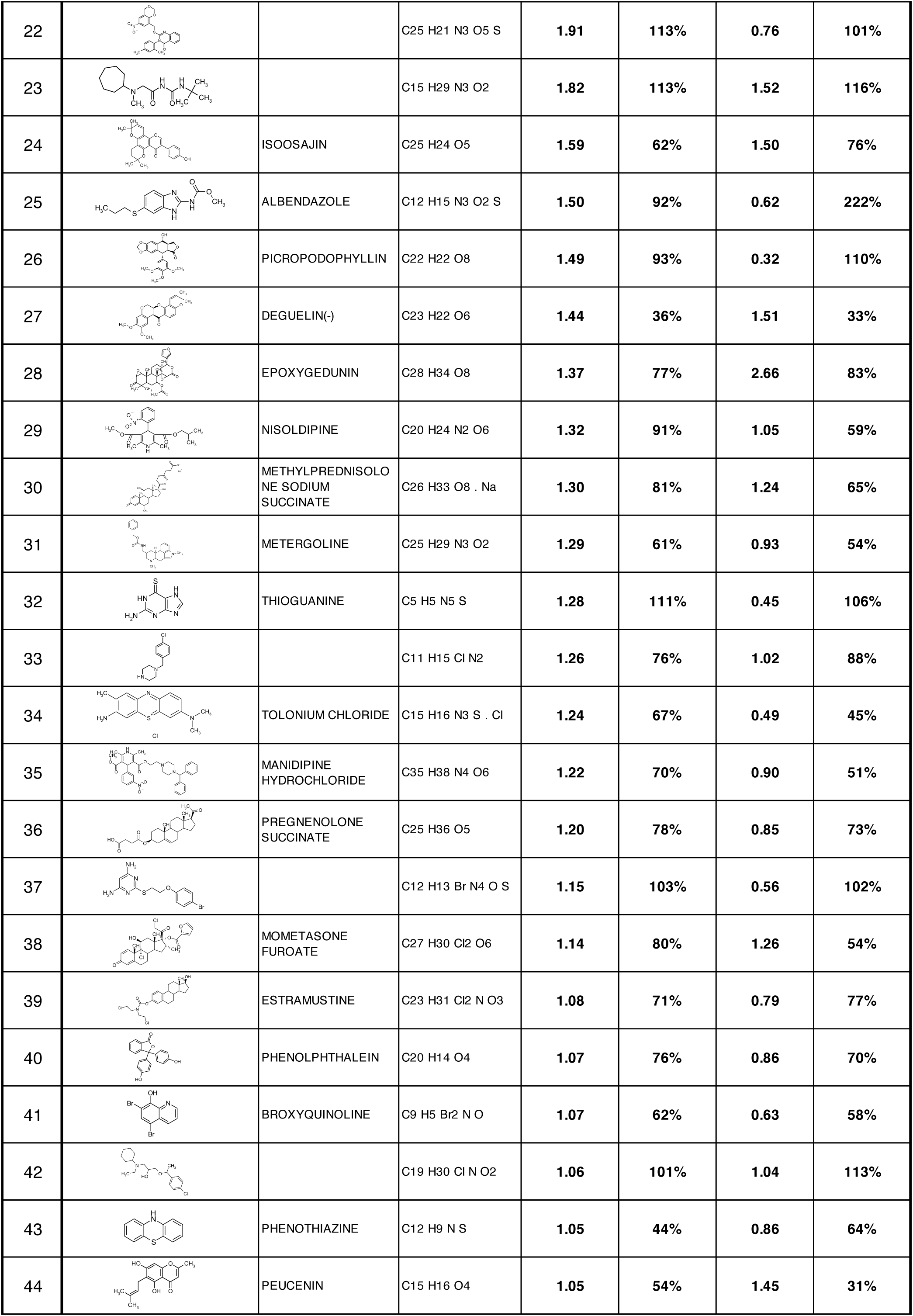
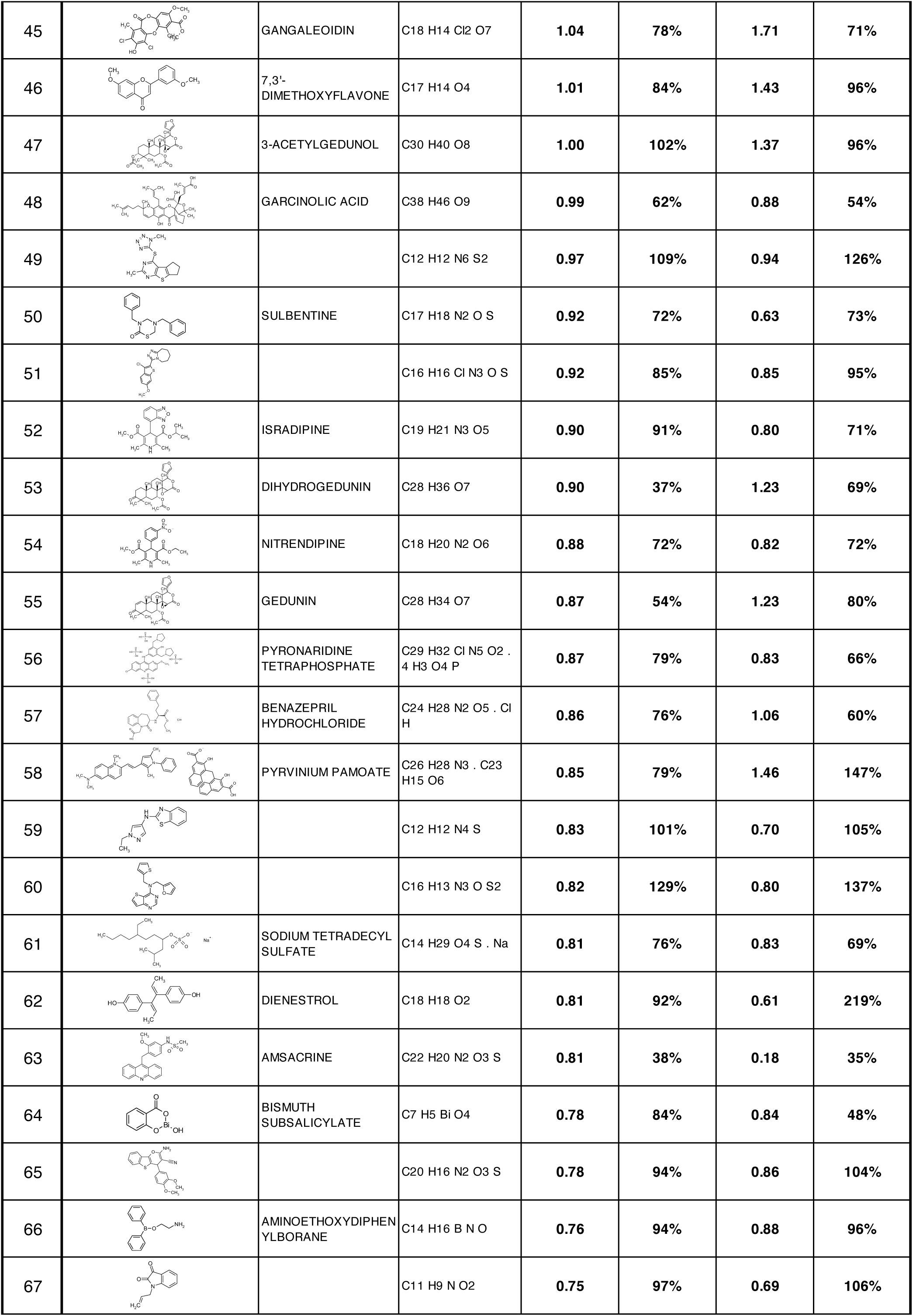
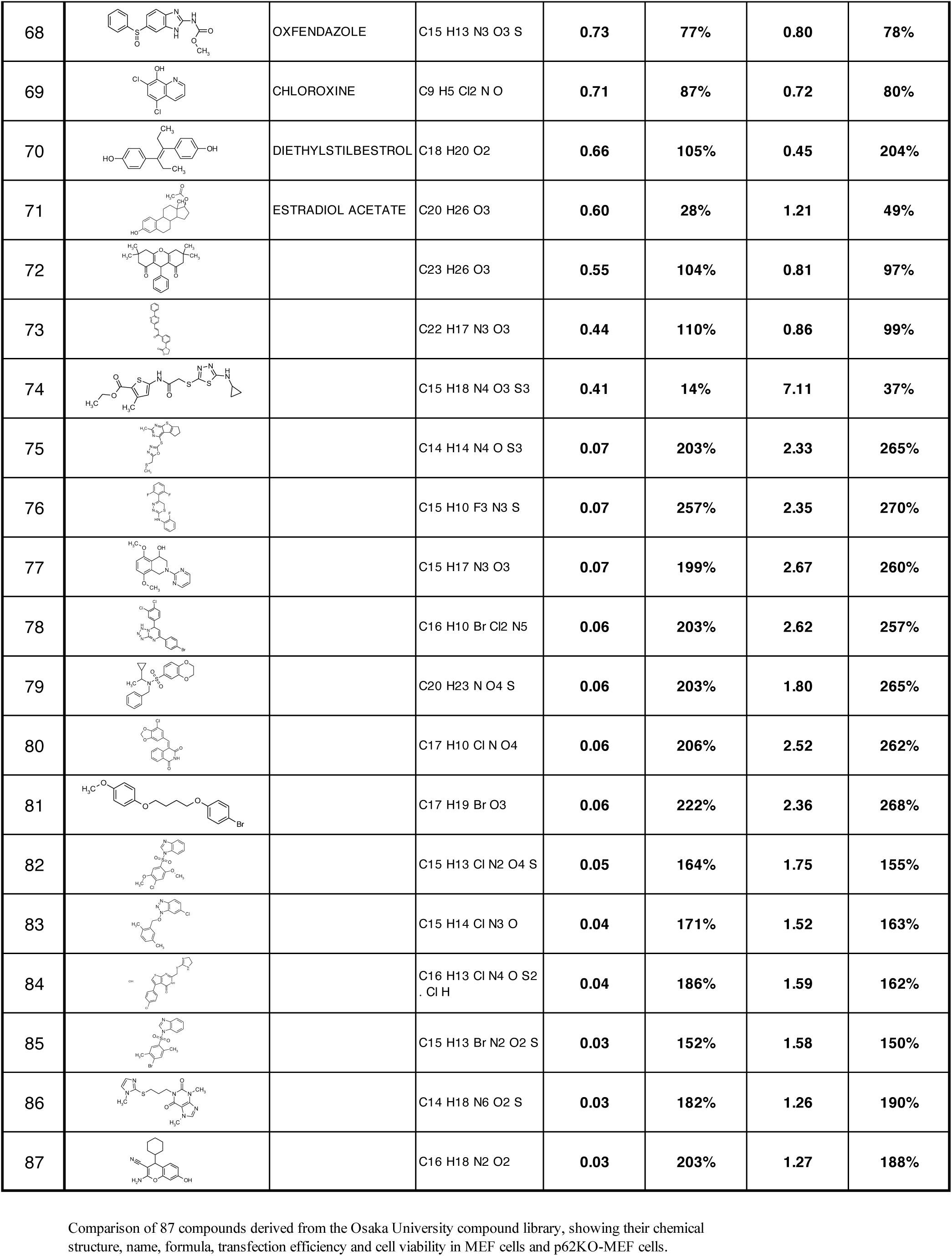
Identification of 87 potential compounds for enhancement of transfection efficiency.

| Rank | Structure | Name | Formula | Fold transfection efficiency in MEF | Fold cell viability in MEF | Fold transfection efficiency in p62KO-MEF | Fold cell viability in p62KO-MEF |
| --- | --- | --- | --- | --- | --- | --- | --- |
| 1 |  | PODOFILOX | C22 H22 O8 | 297.24 | 64% | 4.46 | 281% |
| 2 |  | COLCHICINE | C22 H25 N O6 | 294.20 | 95% | 4.94 | 258% |
| 3 |  | VINBLASTINE SULFATE | C46 H58 N4 O9 . H2 O4 S | 263.07 | 68% | 6.90 | 318% |
| 4 |  | DEOXSAPPANONE B 7,4'-DIMETHYL ETHER | C18 H18 O5 | 54.43 | 96% | 4.72 | 95% |
| 5 |  | VINCRISTINE SULFATE | C46 H56 N4 O10 . H2 O4 S | 46.77 | 40% | 0.66 | 47% |
| 6 |  | PICROPODOPHYLLIN ACETATE | C24 H24 O9 | 35.46 | 34% | 2.19 | 46% |
| 7 |  | DOCETAXEL | C43 H53 N O14 . 3 H2 O | 33.78 | 72% | 9.43 | 316% |
| 8 |  | FENBENDAZOLE | C15 H13 N3 O2 S | 27.14 | 63% | 7.83 | 337% |
| 9 |  | FLUBENDAZOLE | C17 H13 F N2 O3 | 21.35 | 71% | 4.53 | 357% |
| 10 |  |  | C18 H19 Cl F N O2 | 8.77 | 59% | 1.09 | 80% |
| 11 |  |  | C18 H17 N O2 S2 | 6.80 | 74% | 1.80 | 68% |
| 12 |  |  | C15 H26 N4 O5 S2 | 6.19 | 89% | 1.70 | 91% |
| 13 |  | OXIBENDAZOLE | C12 H15 N3 O3 | 4.64 | 45% | 0.85 | 49% |
| 14 |  | DEOXSAPPANONE B 7,3'-DIMETHYL ETHER ACETATE | C20 H20 O6 | 4.50 | 82% | 2.07 | 85% |
| 15 |  | PHENETHYL CAFFEATE (CAPE) | C17 H16 O4 | 4.44 | 72% | 1.63 | 89% |
| 16 |  | MEBENDAZOLE | C16 H13 N3 O3 | 3.01 | 95% | 1.57 | 365% |
| 17 |  | AZACITIDINE | C8 H12 N4 O5 | 2.92 | 59% | 0.93 | 67% |
| 18 |  |  | C15 H10 Br F N2 O2 | 2.84 | 96% | 0.53 | 109% |
| 19 |  |  | C15 H18 N4 O3 | 2.52 | 97% | 0.62 | 105% |
| 20 |  | PIROCTONE OLAMINE | C14 H23 N O2 . C2 H7 N O | 2.04 | 40% | 0.78 | 42% |
| 21 |  |  | C18 H16 N2 | 1.96 | 94% | 0.38 | 102% |
| 22 |  |  | C25 H21 N3 O5 S | 1.91 | 113% | 0.76 | 101% |
| 23 |  |  | C15 H29 N3 O2 | 1.82 | 113% | 1.52 | 116% |
| 24 |  | ISOOSAJIN | C25 H24 O5 | 1.59 | 62% | 1.50 | 76% |
| 25 |  | ALBENDAZOLE | C12 H15 N3 O2 S | 1.50 | 92% | 0.62 | 222% |
| 26 |  | PICROPODOPHYLLIN | C22 H22 O8 | 1.49 | 93% | 0.32 | 110% |
| 27 |  | DEGUELIN(-) | C23 H22 O6 | 1.44 | 36% | 1.51 | 33% |
| 28 |  | EPOXYGEDUNIN | C28 H34 O8 | 1.37 | 77% | 2.66 | 83% |
| 29 |  | NISOLDIPINE | C20 H24 N2 O6 | 1.32 | 91% | 1.05 | 59% |
| 30 |  | METHYLPREDNISOLONE SODIUM SUCCINATE | C26 H33 O8 . Na | 1.30 | 81% | 1.24 | 65% |
| 31 |  | METERGOLINE | C25 H29 N3 O2 | 1.29 | 61% | 0.93 | 54% |
| 32 |  | THIIOGUANINE | C5 H5 N5 S | 1.28 | 111% | 0.45 | 106% |
| 33 |  |  | C11 H15 Cl N2 | 1.26 | 76% | 1.02 | 88% |
| 34 |  | TOLONIUM CHLORIDE | C15 H16 N3 S . Cl | 1.24 | 67% | 0.49 | 45% |
| 35 |  | MANIDIPINE HYDROCHLORIDE | C35 H38 N4 O6 | 1.22 | 70% | 0.90 | 51% |
| 36 |  | PREGNENOLONE SUCCINATE | C25 H36 O5 | 1.20 | 78% | 0.85 | 73% |
| 37 |  |  | C12 H13 Br N4 O S | 1.15 | 103% | 0.56 | 102% |
| 38 |  | MOMETASONE FUROATE | C27 H30 Cl2 O6 | 1.14 | 80% | 1.26 | 54% |
| 39 |  | ESTRAMUSTINE | C23 H31 Cl2 N O3 | 1.08 | 71% | 0.79 | 77% |
| 40 |  | PHENOLPHTHALEIN | C20 H14 O4 | 1.07 | 76% | 0.86 | 70% |
| 41 |  | BROXYQUINOLINE | C9 H5 Br2 N O | 1.07 | 62% | 0.63 | 58% |
| 42 |  |  | C19 H30 Cl N O2 | 1.06 | 101% | 1.04 | 113% |
| 43 |  | PHENOTHIAZINE | C12 H9 N S | 1.05 | 44% | 0.86 | 64% |
| 44 |  | PEUCENIN | C15 H16 O4 | 1.05 | 54% | 1.45 | 31% |
| 45 |  | GANGALEOIDIN | C18 H14 Cl2 O7 | 1.04 | 78% | 1.71 | 71% |
| 46 |  | 7,3'-DIMETHOXYFLAVONE | C17 H14 O4 | 1.01 | 84% | 1.43 | 96% |
| 47 |  | 3-ACETYLGEDUNOL | C30 H40 O8 | 1.00 | 102% | 1.37 | 96% |
| 48 |  | GARCINOLIC ACID | C38 H46 O9 | 0.99 | 62% | 0.88 | 54% |
| 49 |  |  | C12 H12 N6 S2 | 0.97 | 109% | 0.94 | 126% |
| 50 |  | SULBENTINE | C17 H18 N2 O S | 0.92 | 72% | 0.63 | 73% |
| 51 |  |  | C16 H16 Cl N3 O S | 0.92 | 85% | 0.85 | 95% |
| 52 |  | ISRADIPINE | C19 H21 N3 O5 | 0.90 | 91% | 0.80 | 71% |
| 53 |  | DIHYDROGEDUNIN | C28 H36 O7 | 0.90 | 37% | 1.23 | 69% |
| 54 |  | NITRENDIPINE | C18 H20 N2 O6 | 0.88 | 72% | 0.82 | 72% |
| 55 |  | GEDUNIN | C28 H34 O7 | 0.87 | 54% | 1.23 | 80% |
| 56 |  | PYRONARIDINE<br>TETRAPHOSPHATE | C29 H32 Cl N5 O2 .<br>4 H3 O4 P | 0.87 | 79% | 0.83 | 66% |
| 57 |  | BENAZEPRIL<br>HYDROCHLORIDE | C24 H28 N2 O5 . Cl<br>H | 0.86 | 76% | 1.06 | 60% |
| 58 |  | PYRVINIUM PAMOATE | C26 H28 N3 . C23<br>H15 O6 | 0.85 | 79% | 1.46 | 147% |
| 59 |  |  | C12 H12 N4 S | 0.83 | 101% | 0.70 | 105% |
| 60 |  |  | C16 H13 N3 O S2 | 0.82 | 129% | 0.80 | 137% |
| 61 |  | SODIUM TETRADECYL<br>SULFATE | C14 H29 O4 S . Na | 0.81 | 76% | 0.83 | 69% |
| 62 |  | DIENESTROL | C18 H18 O2 | 0.81 | 92% | 0.61 | 219% |
| 63 |  | AMSACRINE | C22 H20 N2 O3 S | 0.81 | 38% | 0.18 | 35% |
| 64 |  | BISMUTH<br>SUBSALICYLATE | C7 H5 Bi O4 | 0.78 | 84% | 0.84 | 48% |
| 65 |  |  | C20 H16 N2 O3 S | 0.78 | 94% | 0.86 | 104% |
| 66 |  | AMINOETHOXYDIPHEN-YLBORANE | C14 H16 B N O | 0.76 | 94% | 0.88 | 96% |
| 67 |  |  | C11 H9 N O2 | 0.75 | 97% | 0.69 | 106% |
| 68 |  | OXFENDAZOLE | C15 H13 N3 O3 S | 0.73 | 77% | 0.80 | 78% |
| 69 |  | CHLOROXINE | C9 H5 Cl2 N O | 0.71 | 87% | 0.72 | 80% |
| 70 |  | DIETHYLSTILBESTROL | C18 H20 O2 | 0.66 | 105% | 0.45 | 204% |
| 71 |  | ESTRADIOL ACETATE | C20 H26 O3 | 0.60 | 28% | 1.21 | 49% |
| 72 |  |  | C23 H26 O3 | 0.55 | 104% | 0.81 | 97% |
| 73 |  |  | C22 H17 N3 O3 | 0.44 | 110% | 0.86 | 99% |
| 74 |  |  | C15 H18 N4 O3 S3 | 0.41 | 14% | 7.11 | 37% |
| 75 |  |  | C14 H14 N4 O S3 | 0.07 | 203% | 2.33 | 265% |
| 76 |  |  | C15 H10 F3 N3 S | 0.07 | 257% | 2.35 | 270% |
| 77 |  |  | C15 H17 N3 O3 | 0.07 | 199% | 2.67 | 260% |
| 78 |  |  | C16 H10 Br Cl2 N5 | 0.06 | 203% | 2.62 | 257% |
| 79 |  |  | C20 H23 N O4 S | 0.06 | 203% | 1.80 | 265% |
| 80 |  |  | C17 H10 Cl N O4 | 0.06 | 206% | 2.52 | 262% |
| 81 |  |  | C17 H19 Br O3 | 0.06 | 222% | 2.36 | 268% |
| 82 |  |  | C15 H13 Cl N2 O4 S | 0.05 | 164% | 1.75 | 155% |
| 83 |  |  | C15 H14 Cl N3 O | 0.04 | 171% | 1.52 | 163% |
| 84 |  |  | C16 H13 Cl N4 O S2<br>· Cl H | 0.04 | 186% | 1.59 | 162% |
| 85 |  |  | C15 H13 Br N2 O2 S | 0.03 | 152% | 1.58 | 150% |
| 86 |  |  | C14 H18 N6 O2 S | 0.03 | 182% | 1.26 | 190% |
| 87 |  |  | C16 H18 N2 O2 | 0.03 | 203% | 1.27 | 188% |

Comparison of 87 compounds derived from the Osaka University compound library, showing their chemical structure, name, formula, transfection efficiency and cell viability in MEF cells and p62KO-MEF cells.

**Fig. S1.**
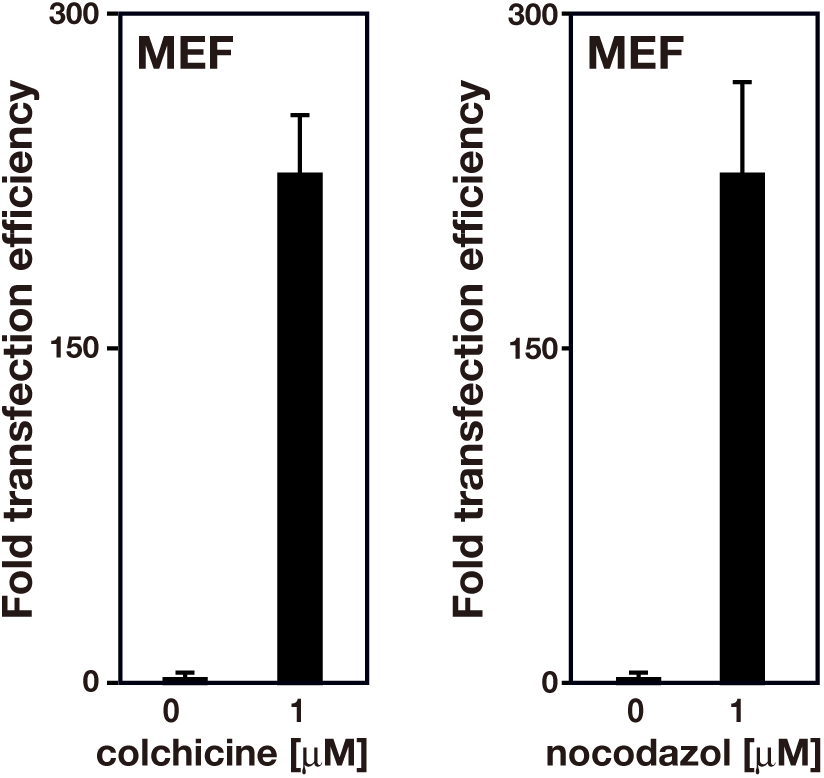
Nocodazole increases gene transfection efficiency. MEF cells were transfected with the pCMV-Luc plasmid for 28 h in the presence of 1 μM colchicine (left) and 1 μM nocodazole (right). DMSO was used as the negative control. Transfection efficiency was determined based on luciferase activity normalized to the cell viability. Bar graphs show the fold changes in transfection efficiency relative to the value from the control DMSO-treated cells. Each value is indicated as the mean ± SD of at least threeindependent experiments.

**Fig. S2.**
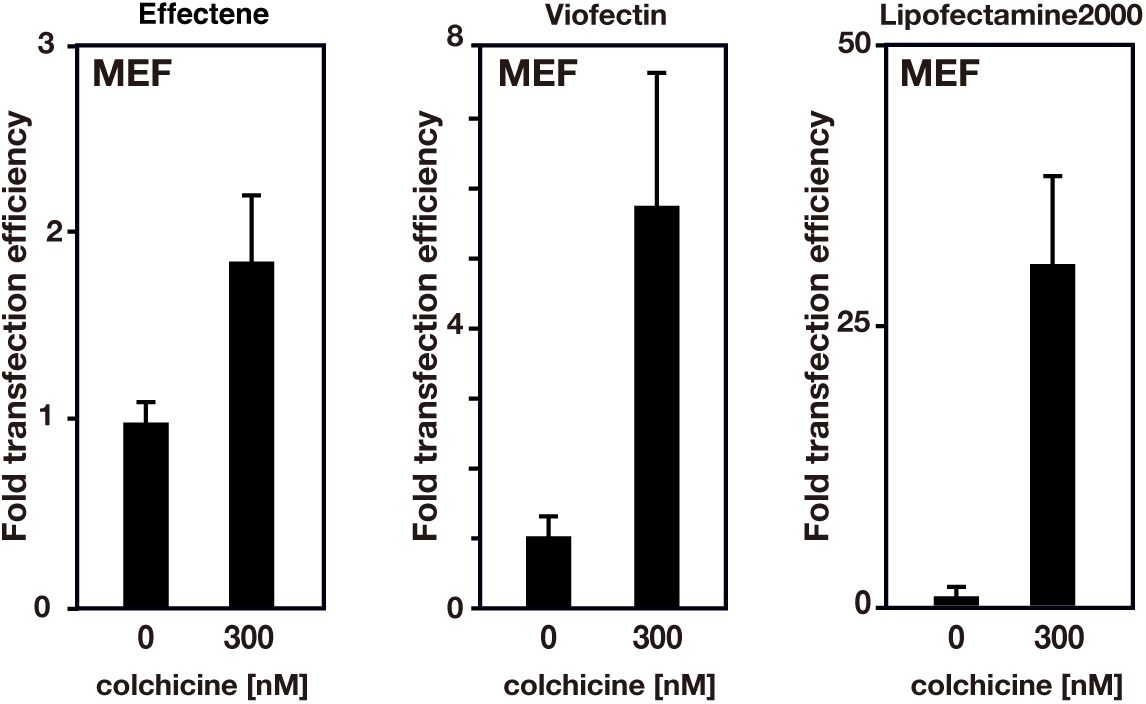
Colchicine increases transfection efficiency for various transfection reagents. MEF cells were transfected with the pCMV-Luc plasmid in the presence of 300 nM colchicine by using Effectene, Viofectin, or Lipofectamine 2000. DMSO was used as the negative control. Details of transfection procedures for each transfection reagent are described in the Experimental procedures. Bar graphs show the fold changes in transfection efficiency relative to the value from the control DMSO-treated cells. (mean ± SD of at least three independent experiments).

**Fig. S3.**
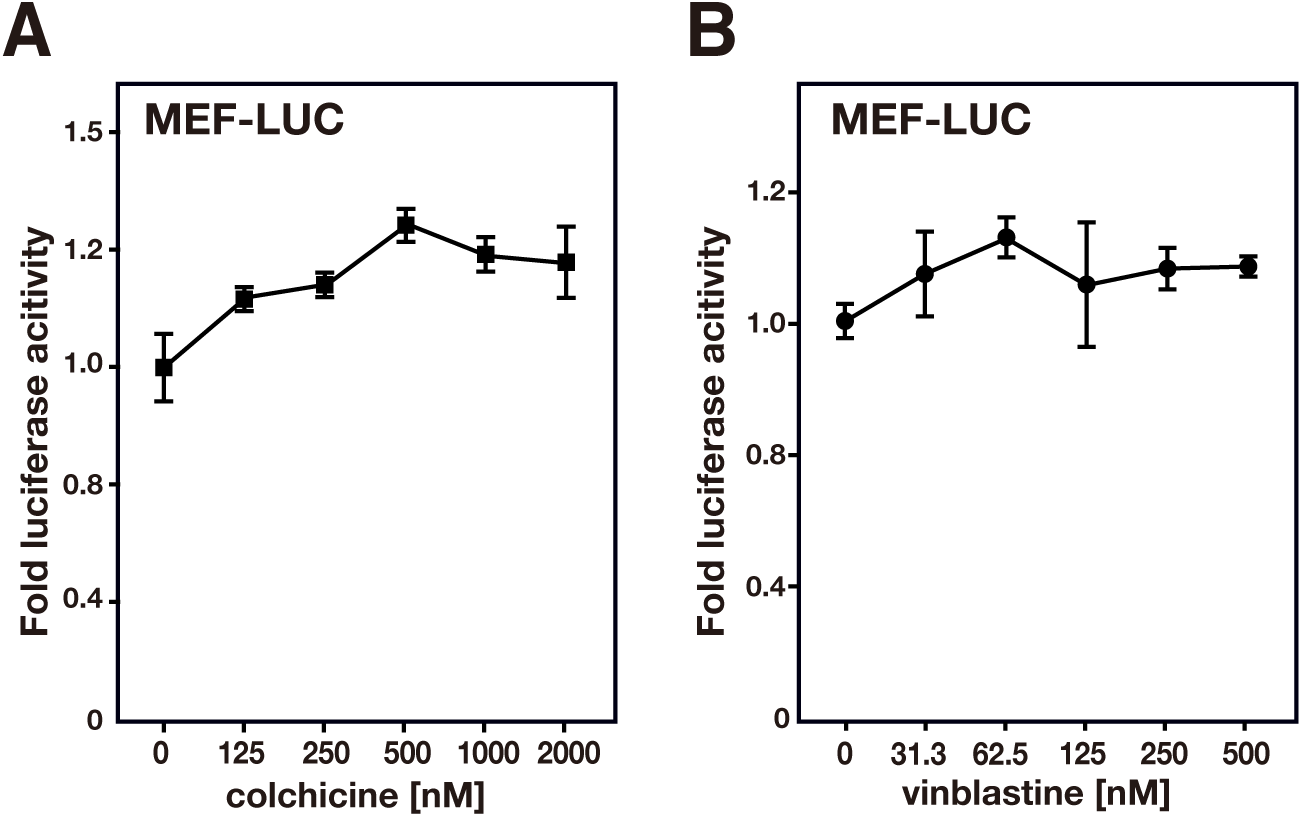
Colchicine and vinblastine treatments do not affect the promoter activity of the luciferase gene. (A, B) MEF cells stably expressing luciferase (MEF-LUC) were treated with the indicated concentrations of colchicine (A) or vinblastine (B). After incubation for 27 h, the luciferase activity of the cells was measured and normalized to cell viability. The graphs show the fold changes in luciferase activity relative to the value from the control DMSO-treated cells (indicated as mean ± SD of at least three independent experimental results).

